# *Saccharomyces cerevisiae* Requires *CFF1* to Produce 4-hydroxy-5-methylfuran-3(2H)-one, a Mimic of the Bacterial Quorum-Sensing Autoinducer AI-2

**DOI:** 10.1101/2020.11.24.397265

**Authors:** Julie S. Valastyan, Christina M. Kraml, Istvan Pelczer, Thomas Ferrante, Bonnie L. Bassler

## Abstract

Quorum sensing is a process of cell-to-cell communication that bacteria use to orchestrate collective behaviors. Quorum sensing depends on the production, release, and detection of extracellular signal molecules called autoinducers (AIs) that accumulate with increasing cell density. While most AIs are species-specific, the AI called AI-2 is produced and detected by diverse bacterial species and it mediates inter-species communication. We recently reported that mammalian cells produce an AI-2 mimic that can be detected by bacteria through the AI-2 receptor, LuxP, potentially expanding the role of the AI-2 system to inter-domain communication. Here, we describe a second molecule capable of inter-domain signaling through LuxP, 4-hydroxy-5-methylfuran-3(2H)-one (MHF) that is produced by the yeast *Saccharomyces cerevisiae*. Screening the *S. cerevisiae* deletion collection revealed Cff1p, a protein with no known role, to be required for MHF production. Cff1p is proposed to be an enzyme, possibly an epimerase or isomerase, and substitution at the putative catalytic residue eliminated MHF production in *S. cerevisiae*. Sequence analysis uncovered Cff1p homologs in many species, primarily bacterial and fungal, but also viral, archaeal, and higher eukaryotic. Cff1p homologs from organisms from all domains can complement a *S. cerevisiae cff1Δ* mutant and restore MHF production. In all test cases, the identified catalytic residue is conserved and required for MHF to be produced. These findings increase the scope of possibilities for inter-domain interactions via AI-2 and AI-2 mimics, highlighting the breadth of molecules and organisms that could participate in quorum sensing.

**Importance:** Quorum sensing is a cell-to-cell communication process that bacteria use to monitor local population density. Quorum sensing relies on extracellular signal molecules called autoinducers (AIs). One AI, called AI-2, is broadly made by bacteria and used for inter-species communication. Here, we describe a eukaryotic AI-2 mimic, 5-methylfuran-3(2H)-one, (MHF), that is made by the yeast *Saccharomyces cerevisiae*, and we identify the Cff1p protein as essential for MHF production. Hundreds of viral, archaeal, bacterial, and eukaryotic organisms possess Cff1p homologs. This finding, combined with our results showing that homologs from all domains can replace *S. cerevisiae* Cff1p, suggests that like AI-2, MHF is widely produced. Our results expand the breadth of organisms that may participate in quorum-sensing-mediated interactions.

## Introduction

Bacteria use chemical communication to gauge local cell population density. This process, called quorum sensing, relies on the production, release, accumulation, and detection of extracellular signal molecules called autoinducers (AIs) (for recent reviews, see (1–3)). Quorum sensing enables bacteria to assess whether they are at low or high cell density and, if the latter, engage in collective behaviors that, to be successful, require many cells acting in synchrony. For example, quorum sensing controls traits such as bioluminescence, biofilm formation, and virulence factor production.

The bioluminescent marine bacterium *Vibrio harveyi* is a model organism used to study quorum sensing. *V. harveyi* employs three AIs, AI-1, CAI-1, and AI-2, that enable intra-species, intra-genera, and inter-species communication, respectively (4–7). Germane to this report is that AI-2 is bound by the receptor LuxP, and LuxP-AI-2 binding initiates a signal transduction cascade, the output of which is bioluminescence (5, 8, 9). AI-2 is produced and detected by diverse bacterial species (6, 10–12). Furthermore, human epithelial cells secrete an AI-2 mimic (here designated “mammalian AI-2 mimic”) of unknown structure that can be detected by LuxP, suggesting that the AI-2 signaling pathway could underpin inter-domain communication (13).

Regarding AI-2 biosynthesis: the AI-2 precursor, 4,5-dihydroxy-2,3-pentanedione (DPD, Figure 1A), is produced by the LuxS synthase from *S*-ribosylhomocysteine (SRH), a biosynthetic intermediate in SAM-dependent methylation pathways (6). DPD, the precursor to all AI-2 moieties, rapidly interconverts between different forms, and these rearranged structures can show preferences for binding to a particular bacterial quorum-sensing receptor. To activate the vibrio LuxP receptor, DPD must cyclize and coordinate borate to form the active AI-2 signaling moiety, *S*-THMF-borate (Figure 1A) (8). The marine environment is borate-rich, favoring formation of this final, borated signal molecule employed by *V. harveyi* and other vibrios. In boron-limited terrestrial environments, DPD rearranges to form *R*-THMF, the active AI-2 moiety detected by enteric bacteria via a LuxP homolog called LsrB (Figure 1A) (14). Another family of receptors, each of which contains a pCACHE domain, was recently discovered, expanding AI-2 detection mechanisms and the breadth of bacterial species that apparently respond to AI-2 (15).

**Figure 1.**
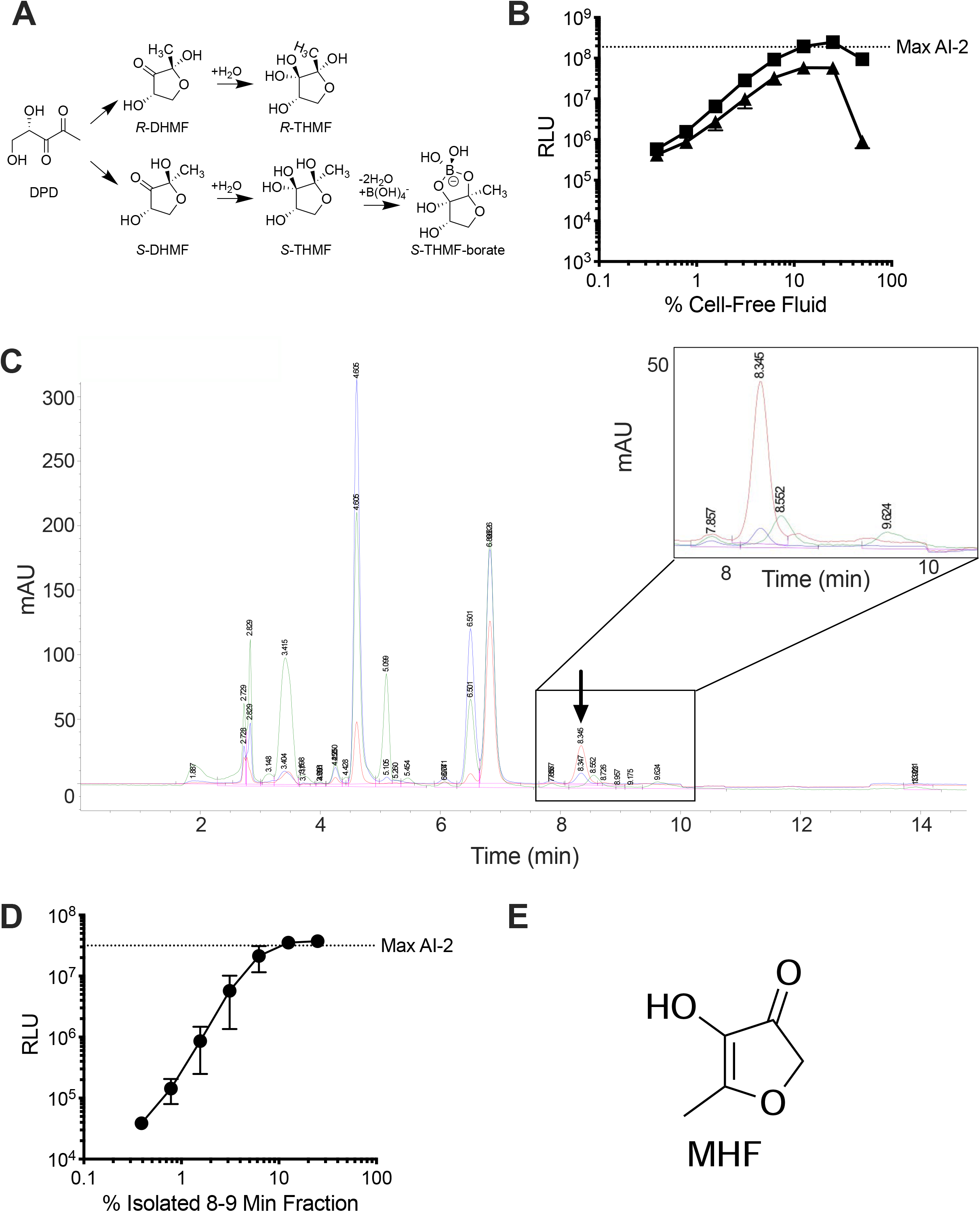
*S. cerevisiae* produces MHF, an AI-2 mimic. (A) Diagram showing the AI-2 biosynthetic pathway from DPD and relevant interconversions among molecules. (B) Light output by the *V. harveyi* TL-26 reporter strain in response to *S. cerevisiae* culture fluids containing yeast AI-2 mimic in PBS (squares) and in water (triangles). (C) Chromatogram depicting fractionation of yeast AI-2 mimic preparations. The area containing the active fraction is enlarged in the inset. The chromatograms show absorption at 214 (green), 254 (blue), and 280 (red) nm. The arrow depicts the peak containing the activity. mAU = milli-Absorbance Units. (D) Light output from the *V. harveyi* TL-26 reporter strain in response to a titration of the active 8-9 min fraction from panel C. (E) Structure of MHF. In B and D, RLU denote relative light units, which are bioluminescence/OD_600_ of the reporter strain and the dotted line labeled Max AI-2 refers to the activity from 125 nM DPD. In B and D, error bars represent standard deviations of biological replicates, N=3.

Fungi also rely on quorum sensing to control behavior. In *S. cerevisiae*, production of phenylethanol and tryptophol drives filamentous growth via activation of expression of *FLO11*, encoding a glycoprotein required for flocculation and biofilm formation (16). *Candida albicans* uses two quorum-sensing molecules to regulate the transition from yeast to filamentous growth: tyrosol promotes the development of germ tubes required for hyphal growth, while farnesol inhibits this transition (17–20). Farnesol also mediates inter-domain interactions by preventing toxin production by the bacterial pathogen *Pseudomonas aeruginosa* when in a mixed population with *C. albicans* (21, 22). Finally, farnesol is reported to modulate the human immune response in a *C. albicans* infection model (23–25). These findings hint at chemically-mediated inter-domain communication between fungi and other prokaryotic and eukaryotic organisms.

Here, we report a new inter-domain quorum-sensing interaction. *S. cerevisiae* produces 4-hydroxy-5-methylfuran-3(2H)-one, a compound that mimics bacterial AI-2. Using *V. harveyi* as a reporter of AI-2 activity, we show that detection of and response to MHF requires the LuxP receptor and signal transduction through the canonical *V. harveyi* quorum-sensing pathway. Screening of the *S. cerevisiae* deletion library revealed Cff1p as a protein essential for MHF production. Cff1p is uncharacterized but its crystal structure shows homology to sugar epimerases and isomerases (26). Mutation of a predicted catalytic residue, glutamic acid at position 44, eliminates MHF production by *S. cerevisiae*, suggesting that Cff1p may function as the MHF synthase. Many putative *CFF1* homologs exist in viral, archaeal, bacterial, fungal, and higher organismal genomes, and in the majority of cases we tested, the *CFF1* genes could complement a *cff1*Δ *S. cerevisiae* mutant and restore MHF production. Alignment of Cff1p homologs showed that the key glutamic acid residue is conserved, and in our test cases, it is required for activity. In summary, MHF has the ability to mimic AI-2, and MHF production may be prevalent in both prokaryotic and eukaryotic organisms. These findings highlight the expanding possibilities for inter-domain signaling through AI-2 quorum-sensing pathways.

## Results

### Purification and identification of 4-hydroxy-5-methylfuran-3(2H)-one (MHF) as an AI-2 mimic produced by S. cerevisiae

We previously reported that human tissue culture cells of epithelial origin, when starved or subjected to tight junction disruption, produce a mimic of the bacterial quorum-sensing AI called AI-2 (13). These earlier findings inspired us to examine whether other eukaryotes produce AI-2 mimics. Here, we focus on the yeast *S. cerevisiae* for two reasons. First, evolutionarily, *S. cerevisiae* and humans diverged ~ 1 billion years ago (27), possibly yielding insight into whether AI-2 mimic production does or does not occur widely across eukaryotes. Second, *S. cerevisiae* can be easily cultured, grows in virtually unlimited quantities, and survives in water, features predicted to accelerate purification and identification of interesting compounds (28). Relevant to this second point is that the identity of the mammalian AI-2 mimic remains unknown, primarily due to the inability to produce sufficient amounts for structural analyses. Moreover, the high salt conditions (PBS) required for mammalian AI-2 mimic production are incompatible with standard purification methods such as HPLC.

We first tested whether *S. cerevisiae* makes a molecule that can mimic AI-2. *S. cerevisiae* MY8092 (hereafter called *S. cerevisiae*) was grown in synthetic defined medium with 2% glucose as the carbon source. Following 48 h of incubation, cell-free culture fluids were prepared and assessed for an activity capable of inducing light production in the *V. harveyi* AI-2 reporter strain called TL-26 (29). *V. harveyi* TL-26 produces maximum light in response to supplementation with 125 nM pure AI-2 (*S*-THMF-borate) (Figure S1A, dotted line). High level activity was present in the *S. cerevisiae* cell-free culture fluids, suggesting that *S. cerevisiae* produces an AI-2 mimic (Figure S1A).

Based on our finding that human epithelial cells produce the mammalian AI-2 mimic when starved in PBS, and, again with the goal of facilitating purification, we next assessed AI-2 mimic production under starvation conditions. *S. cerevisiae* was grown to saturation in rich medium, washed twice, resuspended in either water or PBS and incubated overnight at room temperature. AI-2 mimic activity was present in the collected fluids following resuspension of *S. cerevisiae* in both PBS (Figure 1B, squares) and water (Figure 1B, triangles). Addition of >25% (v/v) of the preparation made in water was toxic to *V. harveyi* TL-26, likely due to the sensitivity of *V. harveyi* to low salt.

To discover whether AI-2 mimic production occurred broadly among wild yeasts or was restricted to laboratory *S. cerevisiae*, a panel of wild *S. cerevisiae* isolates obtained from different environments ranging from clinical settings to vineyards was tested for production of activity using the *V. harveyi* TL-26 reporter (Figure S1B) (30, 31). All the production profiles mirrored that of laboratory *S. cerevisiae,* suggesting that the AI-2 mimic is broadly made by *S. cerevisiae* strains.

To garner sufficient yeast AI-2 mimic for structural analysis, we tested the limit to which we could concentrate the activity. At the final step of the above preparation procedure, the washed *S. cerevisiae* cells were resuspended in water at different cell densities, from OD_600_ = 1 to OD_600_ = 128. Following overnight incubation, cell-free culture fluids were analyzed for activation of light production in *V. harveyi* TL-26. Yeast AI-2 mimic activity increased with increasing *S. cerevisiae* cell density (Figure S1C). Moreover, the activity was specific to the AI-2 quorum-sensing pathway, as light production was not induced by the preparations when supplied to a *V. harveyi* reporter strain (TL-25) that is incapable of detecting AI-2 but, rather, responds exclusively to the *V. harveyi* quorum-sensing AI called AI-1 (3-hydroxy-C4-homoserine lactone) (29, 32) (Figure S1D).

To identify the yeast AI-2 mimic, washed *S. cerevisiae* cells were resuspended in water at OD_600_ = 100. Following overnight incubation, the cell-free fluids were collected and concentrated by lyophilization. HPLC fractionation on a Luna C18 reverse phase column revealed one peak at 8.3 min that exhibited absorption at 254 nm (blue trace, Figure 1C, arrow and inset) and 280 nm (red trace, Figure 1C, arrow and inset). The material did not absorb significantly at 214 nm (Figure 1C inset, green trace). The peak contained high levels of yeast AI-2 mimic activity as judged by the *V. harveyi* TL-26 reporter strain (Figure 1D). We pooled this peak from multiple such column runs and prepared the sample for NMR and mass spectral analyses as described in the Methods.

Identification of the bioactive molecule relied on comparison of results from LC-MS, NMR, and GC-MS. LC-MS analysis showed that two components were present in the active fraction. From our initial HPLC fractionation, these components correspond to the peak with absorption at 254 nm and 280 nm (the active peak) and the peak with absorption at 214 nm (an inactive contaminant), which could not be completely separated for LC-MS. The bioactive component had an exact mass of 115.039 (M+H) and a putative molecular formula of C_5_H_6_O_3_. These data, combined with analysis of key peaks in the ^13^C NMR spectra (signals at δ 194, 172, and 134 ppm), led us to consider structures **A-D**(Figure S2A). Definitive evidence for structure **A** was obtained by GC-MS analysis, including matching of the fragmentation pattern of the active component against a database of known structures (Figure S2B). The yeast AI-2 mimic structure **A** was identified as 4-hydroxy-5-methylfuran-3(2H)-one (MHF, Figure 1E, S2A). Indeed, comparison of mass spectral, NMR, and HPLC analytical data confirmed that MHF purified from *S. cerevisiae* was identical to an authentic commercial sample of MHF.

### The mammalian AI-2 mimic is not MHF

With the MHF structure in hand, we could investigate whether the *S. cerevisiae* and the previously reported mammalian AI-2 mimic are identical or not. As noted earlier, the mammalian AI-2 mimic has not been identified so we do not have purified compound (13). Rather, we made a preparation from Caco-2 cells containing high level mammalian AI-2 mimic activity in PBS (13). To determine the elution pattern for MHF in such Caco-2 cell preparations, we spiked commercial MHF into the mammalian AI-2 mimic preparation prior to HPLC fractionation. MHF eluted at 14 min (Figure S3A, arrow) in the context of Caco-2 culture fluids. Samples from Caco-2 cells that had not been spiked did not have a peak at the expected elution time for MHF (Figure S3B, arrow). To eliminate the possibility that MHF was present in the Caco-2 cell preparations but at a level below the UV detection limit on the HPLC instrument, we tested all of the fractions for activity in the *V. harveyi* TL-26 reporter assay. While the reporter assay showed that the mammalian AI-2 mimic was indeed present in the non-MHF-spiked Caco-2 preparations (Figure S3C, black), there was no activity in the 12-14 min HPLC fraction (Figure S3C, red). Collectively, these data demonstrate that the mammalian AI-2 mimic is not MHF. In future studies, we will focus on identification of the mammalian AI-2 mimic.

### MHF agonizes LuxP with a Nanomolar EC_50_

We next assessed the quantity of MHF present in *S. cerevisiae* cell-free fluids. To do this, we grew *S. cerevisiae*, pelleted and washed the cells, and then resuspended the cells at three different cell densities in water. Following overnight incubation, we removed the cells and compared the activities in each of the cell-free fluids to known quantities of the commercial MHF standard. We used two different MHF quantitation methods (Figure 2A). First, different concentrations of commercial MHF were assessed by HPLC, and the areas under the MHF peaks were used to generate a standard curve. The *S. cerevisiae* preparations were likewise subjected to HPLC analysis and the amount of MHF present in each sample was calculated by interpolating the area under the HPLC peak to that from the standard curve (Figure 2A, black bars). Second, commercial MHF was assessed in the *V. harveyi* TL-26 reporter assay at different concentrations to generate an activity-based standard curve. The *S. cerevisiae* cell-free fluids were identically assayed, and the MHF concentration in each preparation was estimated from the activity standard curve (Figure 2A, white bars). The concentrations of MHF in the preparations calculated by the two methods were in close agreement. Assuming MHF production has a linear relationship with OD_600_, we can use our data to estimate that *S. cerevisiae* produced 1.2 ± 0.4 μM MHF per OD_600_ of cells. In the context of detection by the *V. harveyi* quorum-sensing apparatus, the EC_50_ for AI-2 is 3 nM and that for MHF is 300 nM (Figure 2B). Thus, while both compounds exert activity in this system within the range reported for bacterial AIs (33–36), the LuxP receptor prefers AI-2 over MHF.

**Figure 2.**
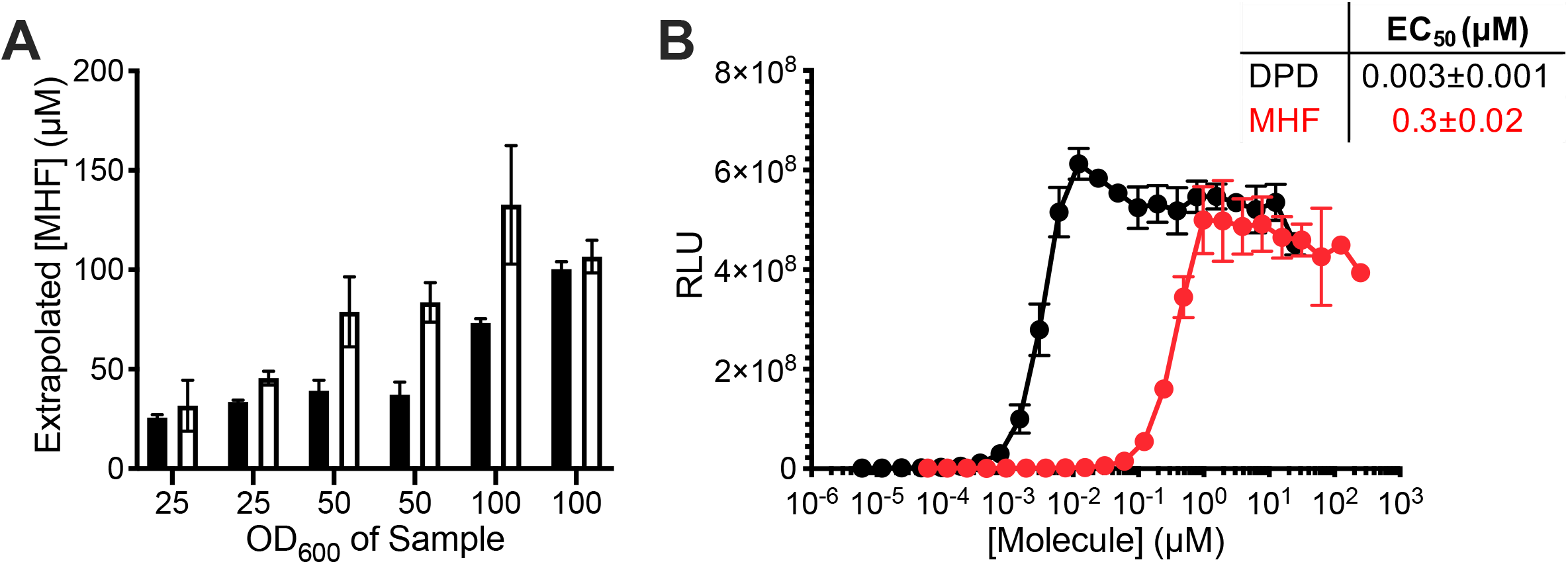
MHF agonizes the LuxP receptor with a nanomolar EC_50_. (A) Quantitation of MHF levels in yeast AI-2 mimic preparations from *S. cerevisiae* concentrated to OD_600_ = 25, 50, or 100 using integration under HPLC peaks (black) or activity from the *V. harveyi* TL-26 reporter strain (white). (B) Light output by the *V. harveyi* TL-26 reporter strain in response to DPD (black) or MHF (red). The table shows the EC_50_ values. RLU as in Figure 1. In A, error bars represent standard deviations of technical replicates, N=3. In B, error bars represent standard deviations of biological replicates, N=3.

### Identification of CFF1 as an S. cerevisiae gene essential for MHF production

To identify the component(s) responsible for MHF production in *S. cerevisiae*, we screened the yeast deletion library for an *S. cerevisiae* mutant that was defective in MHF production (37–39). As described in the Methods, cell-free fluid preparations were made from >5,000 *S. cerevisiae* mutants and incubated with the *V. harveyi* TL-26 reporter strain. Bioluminescence was measured to assess the ability of each *S. cerevisiae* mutant to make MHF (Figure 3A). Mutants were identified that elicited at least two standard deviations less light from the reporter strain than the mean amount of light production elicited from all strains (Figure 3A). Eight putative mutants were retested for the ability to make activity (Figure 3B). Two mutants, *cff1Δ* and *rps1bΔ* failed to activate the reporter strain. Cff1p (systematic name – YML079wp) is a cupin superfamily protein (26). Rps1bp (systematic name – YML063wp) is a component of the 40S ribosomal subunit (40). The six other potential mutants proved to have been false positives upon reassessment (Figure 3B).

**Figure 3.**
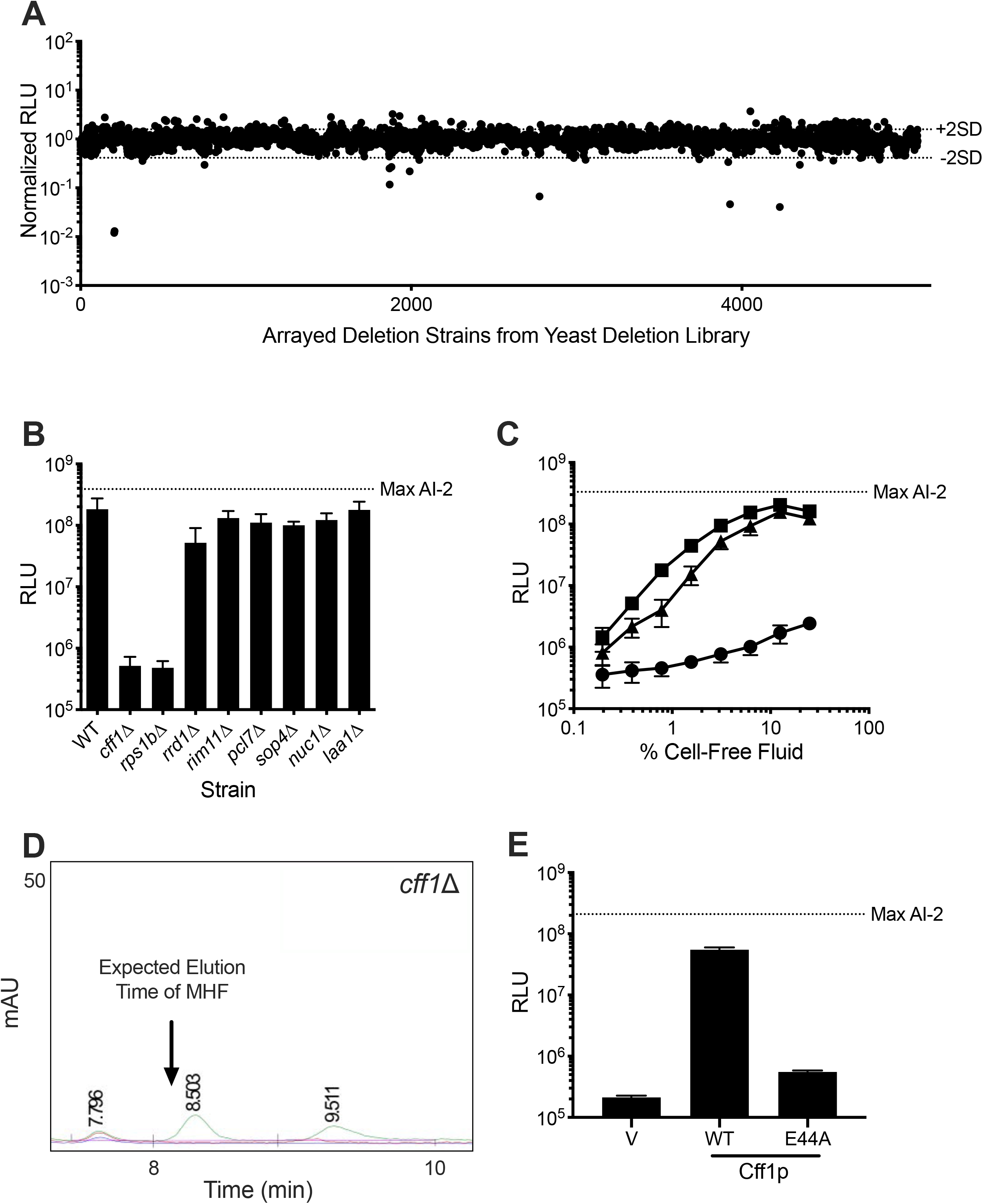
Cff1p is required for *S. cerevisiae* to produce MHF. (A) Normalized light output from the *V. harveyi* TL-26 reporter strain in response to culture fluids from the mutants in the *S. cerevisiae* deletion library. Each point represents the reporter response to a fluid made from a unique yeast mutant. Dotted lines labeled + 2SD and – 2SD show two standard deviations above and below the mean, respectively. (B) Light output from the *V. harveyi* TL-26 reporter strain in response to culture fluids from the putative hit *S. cerevisiae* mutants from panel A. (C) Light output from the *V. harveyi* TL-26 reporter strain in response to culture fluids from WT *S. cerevisiae* (squares) and the *cff1Δ* (circles) and *rps1bΔ* (triangles) mutants. (D) Portion of an HPLC trace from fractionation of yeast AI-2 mimic preparation made from *S. cerevisiae cff1Δ*. The chromatograms show absorption at 214 (green), 254 (blue), and 280 (red) nm. The arrow shows the expected elution time for MHF based on WT *S. cerevisiae* results (see Figure 1C). (E) Light output from the *V. harveyi* TL-26 reporter strain in response to cell-free fluids made from *cff1Δ S. cerevisiae* that produced either a HALO control (designated “V”), Cff1p-HALO (designated WT), or Cff1p-E44A-HALO (designated E44A). 10% (v/v) of cell-free fluid was added in each case. Normalized RLU in A are RLU of the given sample divided by the average RLU from all plates assayed on a single day. In B, C, and E, RLU and Max AI-2 as in Figure 1. In B, C, and E, error bars represent standard deviations of biological replicates, N=3.

To verify the phenotypes of the mutants, clean deletions of *CFF1* and *RPS1B* were constructed in *S. cerevisiae*. The *cff1*Δ mutant displayed no defect in growth rate (Figure S4A, circles; compare to WT growth shown by the squares). As previously reported (41), the *rps1bΔ* mutant had a growth defect (Figure S4A, triangles). Neither mutant exhibited sensitivity to overnight incubation in water (Figure S4B). Importantly, culture fluids prepared from the clean *rps1bΔ* mutant produced nearly the wildtype level of AI-2 mimic activity, as determined by the ability to induce light production in the *V. harveyi* TL-26 reporter (Figure 3C, triangles). By contrast, preparations made from the *cff1Δ* mutant had no activity (Figure 3C, circles). PCR analysis revealed that the mutant annotated as *rps1bΔ* in the yeast deletion library, in fact, possesses a deletion in *CFF1*, explaining its inability to stimulate the reporter strain as well as the ability of our newly constructed *rps1bΔ* mutant to produce activity. Thus, *CFF1* is the only gene revealed by our screen to be required for production of the activity we are monitoring.

To confirm that the yeast AI-2 mimic activity produced by the protein encoded by *CFF1* is MHF, we prepared and fractionated cell-free culture fluids from the *cff1Δ* strain using the identical procedure we used for isolation of MHF from wildtype *S. cerevisiae*. No MHF peak could be detected in the *cff1Δ* mutant preparation (Figure 3D). Consistent with this finding, the relevant HPLC column fraction had no activity in the *V. harveyi* TL-26 reporter assay (Figure S4C). These data suggest that Cff1p has a required role in MHF biosynthesis in yeast.

Our finding that Cff1p is required for MHF production is surprising. The presence of MHF in fermented food products made from *S. cerevisiae* has been reported; however, the suggested route to MHF is either spontaneous starting from D-ribulose-5-phosphate (42–45) or under extreme conditions, via the Maillard reaction (46, 47). Quite to the contrary, our data suggest that MHF production in *S. cerevisiae* is enzyme-catalyzed and under physiological conditions. Cff1p has not been characterized. However, there does exist a crystal structure (26). It shows a putative ligand binding pocket containing amino acid residues identical to those required for catalysis by epimerases and isomerases that share the cupin fold (48). Specifically, the conserved E44 residue is proposed to have a catalytic role. We made an E44A substitution in Cff1p and assayed the mutant protein for MHF production. The substitution did not alter Cff1p stability as judged by visualization of a fused HALO tag (Figure S4D); however, culture fluids prepared from the *S. cerevisiae* Cff1p-E44A mutant elicited 100-fold less light from the *V. harveyi* TL-26 reporter strain than did preparations from WT *S. cerevisiae* (i.e., less than 1% activity remained, Figure 3E) showing that the glutamate residue at position 44 is key for the presumptive enzymatic activity that generates MHF.

### CFF1 homologs exist in organisms from all domains

Cff1p has structural similarity to sugar isomerases and epimerases, and the DNA encoding *CFF1* is similar to genes of unknown function (26). Recently, Tourneroche *et al.* reported twelve wild fungal species that exist as endomicrobiota of kelp and that possess AI-2 activity as judged by a *V. harveyi* reporter system analogous to the one we use here (49). The molecule(s) responsible for the AI-2 activity have not been identified. We wondered whether these fungal species might make MHF. Examination of their proteomes revealed Cff1p homologs in 2 of the 12 species, *Trametes versicolor* and *Botrytis cinerea*, with, respectively, approximately 50% similarity and 35% identity to *S. cerevisiae* Cff1p at the amino acid sequence level (Figure S5A). Both of these species’ Cff1p homologs have high conservation in the putative ligand binding domain, and they each possess a residue equivalent to E44 in *S. cerevisiae* Cff1p (Figure S5A, arrow). To test for function, we cloned these two genes and introduced them into our *S. cerevisiae cff1Δ* mutant under the control of the native *S. cerevisiae CFF1* promoter. Unlike the *S. cerevisiae cff1Δ* mutant that produced no MHF, the mutant carrying each homolog produced activity sufficient to induce maximal light production in the *V. harveyi* TL-26 reporter strain (Figure 4A). The E38A and E30A substitutions in the, respectively, *B. cinerea* and *T. versicolor* Cff1p proteins (equivalent to E44A in *S. cerevisiae* Cff1p) eliminated production of the activity (Figure 4B). In both cases, the WT and mutant proteins were made at approximately the same levels and were equally stable (Figure S5B). HPLC fractionation confirmed that MHF was indeed produced in *cff1Δ S. cerevisiae* carrying the WT *CFF1* homologs, and no MHF could be detected in the cases in which mutant *CFF1* alleles were present (Figure S6). Thus, both the *T. versicolor* and *B. cinerea CFF1* genes can complement the *cff1Δ S. cerevisiae* defect and restore MHF production, and a glutamate at a position equivalent to 44 in *S. cerevisiae* Cff1p is required. In the cases of the other 10 fungi that Tourneroche *et al.* reported to possess AI-2 activity (49), we do not know whether they make a different active molecule or, alternatively, if they possess Cff1p proteins that are unrecognizable through the database search we performed.

**Figure 4.**
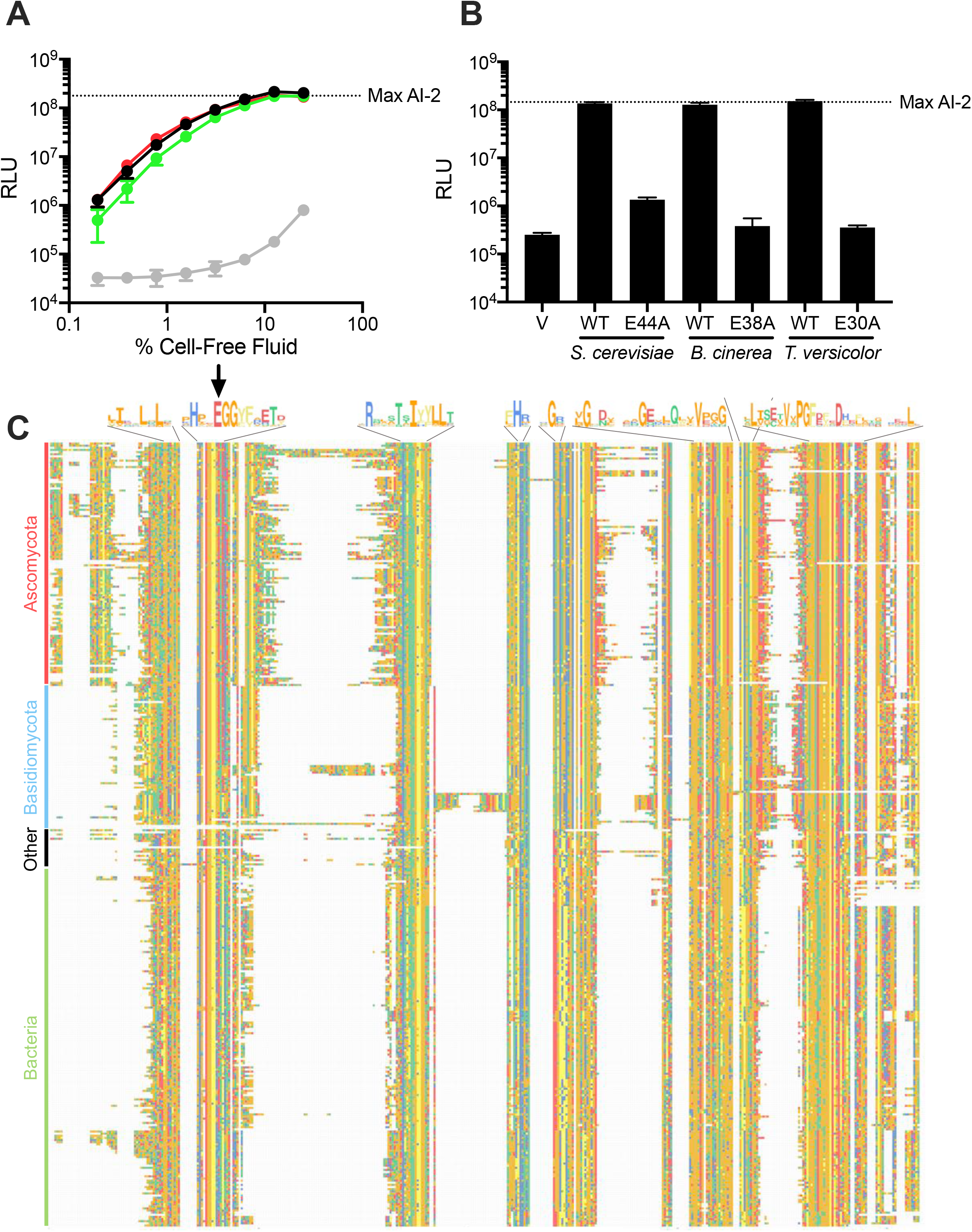
Cff1p homologs restore MHF production to *cff1Δ S. cerevisiae*. (A) Light output from the *V. harveyi* TL-26 reporter strain in response to cell-free fluids from *cff1Δ S. cerevisiae* expressing *CFF1* homologs from *B. cinerea* (green), *T. versicolor* (red), *S. cerevisiae* (black), and a vector control (gray). (B) Light output from the *V. harveyi* TL-26 reporter strain in response to cell-free fluids prepared from *cff1Δ S. cerevisiae* carrying the designated *CFF1* homologs and alleles. The vector control is designated “V”. Cell-free fluids were added at 10% (v/v). (C) Alignment of putative Cff1p homologs, trimmed to the first and final amino acids of *S. cerevisiae* Cff1p. Only one species per genus is shown. Alignment was performed using Clustal Omega (78). Negatively charged residues are shown in red, small hydrophobic residues in orange, aromatic hydrophobic residues in yellow, polar uncharged residues in green, and positively charged residues in blue. Shown above the alignment are consensus sequences for regions containing amino acids that are conserved in >75% of the aligned proteins. Arrow designates location of conserved glutamate residue. RLU and Max AI-2 as in Figure 1. In A and B, error bars represent standard deviations of biological replicates, N=3.

A search of the non-redundant protein sequences database (50) for those with homology to *S. cerevisiae*, *T. versicolor*, and *B. cinerea* Cff1p uncovered three additional *Saccharomyces* species possessing proteins with an average identity of 90% to Cff1p. More broadly, we identified 410 non-*Saccharomyces* fungal species possessing proteins harboring 26%-71% identity to Cff1p. Included were members of the Ascomycota phylum, such as *Aspergillus fumigatus* and *Neurospora crassa*, and the Basidiomycota phyla, such as *Cryptococcus neoformans*. We also identified >350 prokaryotes possessing putative proteins with 25%-45% identity to Cff1p. These species exist in 11 phyla and include multiple *Pseudomonads, Staphylococci,* and *Bacilli*. Our search also uncovered 25 other organisms spanning all domains, with potential Cff1p homologs, the majority of which exist in the marine environment, for example, the acorn worm *Saccoglossus kowalevskii*, the green alga *Ostreococcus lucimarinus*, and the archaeon *Methanohalophilus halophilus* (Figures 4C and S7). At present, we do not know whether this representation indicates a dominant biological function in the marine niche or if it stems from biased sampling. Notably, E44 is conserved in 97% of these potential homologs (Figure 4C, arrow).

It is intriguing that bacterial species may possess *cff* genes (note: we designate the bacterial genes and proteins as *cff* and Cff, respectively, and the fungal genes and proteins *CFF1* and Cff1p, respectively). Thousands of bacterial species are known to synthesize the inter-species quorum-sensing AI AI-2 via possession of *luxS* encoding the AI-2 synthase (10). To investigate whether bacteria could potentially make both MHF and AI-2, we performed database analyses. According to the non-redundant protein sequences database (50), only ~20% of bacterial species possessing a *cff* homolog also harbor *luxS* (Figure 5A). The majority of bacteria that possess both *luxS* and *cff* genes belong to the Firmicutes phylum (Figure 5B), a phylum considered ancestral to other bacterial phyla. This pattern suggests that a shared ancestor possessed both genes, and generally, species maintained only one of the two genes as they diverged.

**Figure 5.**
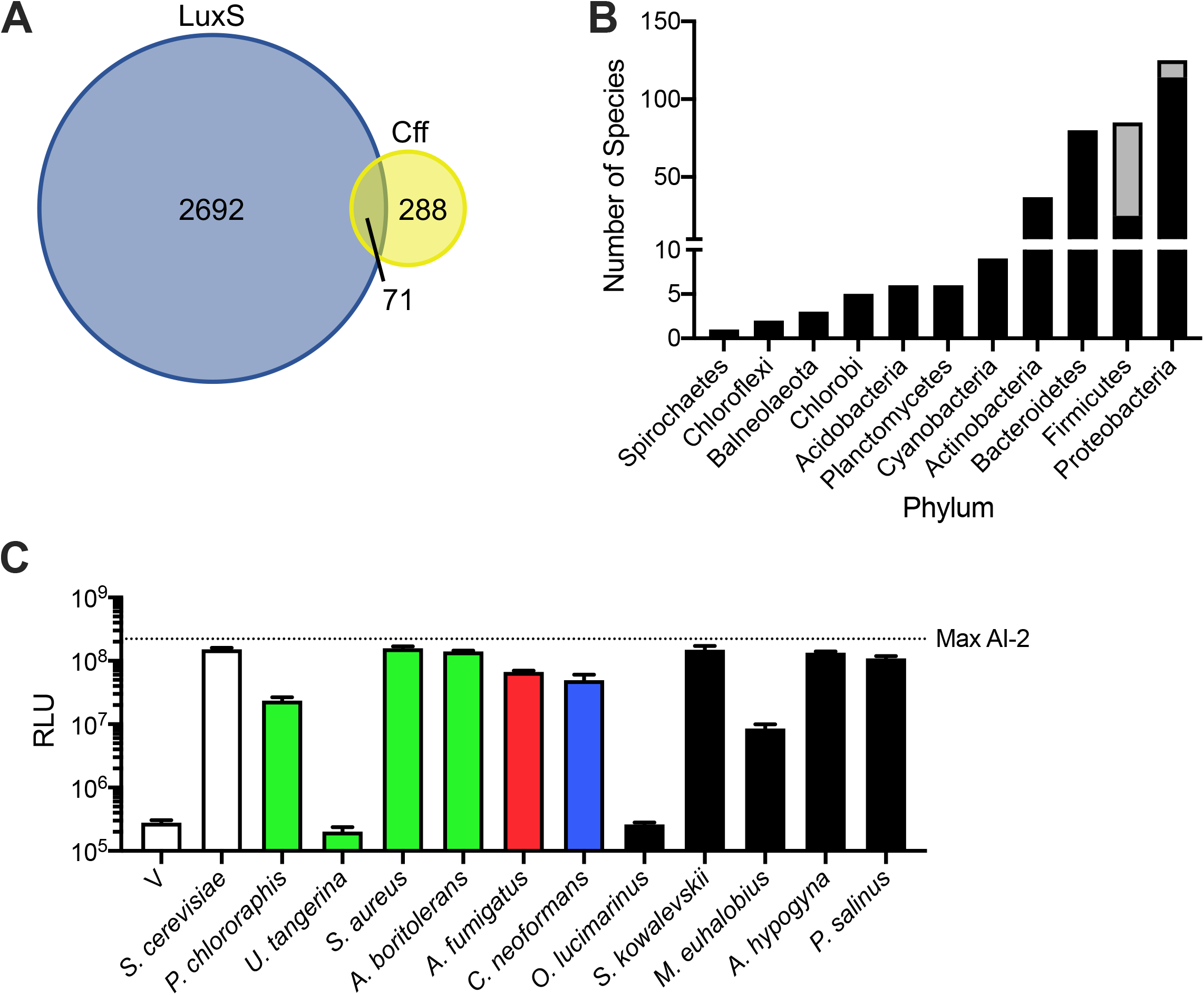
Organisms from all domains contain Cff1p homologs. (A) Venn diagram displaying the numbers of bacterial species containing LuxS and/or Cff homologs. (B) Bacterial phylogeny distribution showing species containing potential Cff homologs. Black bars represent species possessing only Cff. Gray bars represent species possessing both LuxS and Cff. (C) Light output from the *V. harveyi* TL-26 reporter strain in response to cell-free fluids prepared from *S. cerevisiae* expressing *CFF1* and *cff* homologs. Bars are colored according to the groups in Figures 4C and S7: green, Bacteria; red, Ascomycota; blue, Basidiomycota; black, Other. The vector control is designated “V”. Cell-free fluids were added at 10% (v/v). RLU and Max AI-2 as in Figure 1. In C, error bars represent standard deviations of biological replicates, N=3.

To test whether additional Cff and Cff1p homologs are functional, we selected 10 additional organisms – four prokaryotes (*Pseudomonas chlororaphis, Umezawaea tangerina, Staphylococcus aureus,* and *Algoriphagus boritolerans*), two additional fungi (*Aspergillus fumigatus* and *Cryptococcus neoformans*), and four organisms from other domains (*Ostreococcus lucimarinus, Saccoglossus kowalevskii, Methanohalophilus euhalobius,* and *Achlya hypogyna*) that possess putative Cff or Cff1p proteins (Table 1). Our choices included clinically relevant organisms (i.e., *S. aureus* and *C. neoformans*) and marine organisms (i.e., *O. lucimarinus* and *S. kowalevskii*) that may coexist with bacteria that use AI-2-LuxP-mediated quorum sensing. Additionally, we tested proteins from organisms representing unique phyla and proteins with varying levels of amino acid identity relative to *S. cerevisiae* Cff1p (Table 1). To assay activity, we expressed the candidate genes in our *cff1Δ S. cerevisiae* strain under control of the endogenous *S. cerevisiae CFF1* promoter. All of the homologs except those from *U. tangerina* and *O. lucimarinus* complemented the loss of Cff1p in *S. cerevisiae*, driving sufficient MHF production to induce light production in the *V. harveyi* TL-26 reporter strain (Figure 5C). This result suggests that these, and likely many other organisms harbor the potential to synthesize MHF.

**Table 1.** Cff1p homology in select species.

| Phylum | Species | % Coverage | % Identity |
| --- | --- | --- | --- |
| *Bacteria* |  |  |  |
| Proteobacteria | *Pseudomonas chlororaphis* | 76 | 43 |
| Actinobacteria | *Umezawaea tangerina* | 77 | 37 |
| Firmicutes | *Staphylococcus aureus* | 73 | 32 |
| Bacteroides | *Algoriphagus boritolerans* | 77 | 28 |
| *Fungi* |  |  |  |
| Ascomycota | *Botrytis cinerea* | 89 | 44 |
| Basidiomycota | *Trametes versicolor* | 76 | 42 |
| Ascomycota | *Aspergillus fumigatus* | 93 | 40 |
| Basidiomycota | *Cryptococcus neoformans* | 87 | 35 |
| *Other* |  |  |  |
| Algae | *Ostreococcus lucimarinus* | 68 | 41 |
| Animal | *Saccoglossus kowalevskii* | 79 | 38 |
| Archaea | *Methanohalophilus euhalobius* | 76 | 32 |
| Protista | *Achlya hypogyna* | 69 | 31 |
| Virus | *Pandoravirus salinus* | 69 | 24 |

Compared to BLASTp, Interpro (51) provides a dramatically larger set of proteins under the class “Uncharacterized Protein, YML079W-like” (i.e., the *S. cerevisiae* deletion library systematic name for Cff1p). This group contains proteins from >3000 different organisms, including >2400 species of bacteria. 95% of these putative Cff homologs have the conserved glutamate at the position corresponding to 44 in the *S. cerevisiae* Cff1p. The Interpro data suggest that, potentially, the number of bacterial species containing *cff* is on the same scale as those possessing *luxS*. In this dataset, ~15% of the bacterial species contain both *luxS* and *cff*.

We were especially intrigued that the Interpro database search revealed a virus with a potential Cff homolog, *Pandoravirus salinus*. Using the above strategy, we tested the functionality of the *P. salinus* Cff homolog and found that, indeed, the viral *cff* gene complements the loss of *CFF1* in *S. cerevisiae* (Figure 5C). This finding provides initial validation for the output of the larger Interpro dataset and presages the existence of many additional functional Cff1p homologs. Moreover, this final result, coupled with the other findings here, shows that MHF production could occur across the viral, archaeal, bacterial, and eukaryotic domains.

## Discussion

Quorum sensing is the process by which bacteria monitor their local cell population density and determine when it is appropriate to engage in collective behaviors (1–3). Bacteria often employ multiple AIs, encoding distinct information about species relatedness, which presumably enables them to take a census of “self” and “other”. The AI-2-LuxP quorum-sensing pathway is proposed to be used for the latter, to monitor the total cell density of the vicinal community. Previously, we showed that mammalian cells can make a mimic of AI-2 that activates quorum sensing via LuxP, providing a founding example of inter-domain quorum-sensing-mediated communication (13). Here, we show that *S. cerevisiae*, another eukaryotic organism, makes MHF, which can induce quorum-sensing behavior through the canonical AI-2 pathway. Our preliminary studies suggest that MHF production may be widespread across domains. Presumably, the ability of MHF to substitute for AI-2 enables bacteria that possess LuxP to detect the presence of other bacterial cells, higher organisms, and possibly viruses in the environment.

The mammalian AI-2 mimic is distinct from MHF, suggesting that multiple compounds might be exploited for AI-2-like interactions between eukaryotes and bacteria. Plants, insects, and fungi have previously been shown to produce MHF (42–44, 52–54), and based on the findings presented here, it is possible that MHF could be widely used for cross-domain interactions between bacteria and fungi. While future studies are necessary to understand the ecological significance of MHF in such presumptive inter-domain interactions, and, more specifically, what, if any, traits are controlled by MHF in organisms that produce MHF, it is already known that cross-communication between eukaryotes and prokaryotes can shape each participant’s biology. For example, quorum-sensing pathways drive behavior across domains in both mutualistic and parasitic relationships. Examples include communication between the gut and the microbiome (55), competition between *C. albicans* and *P. aeruginosa* during infection of the lung (56), and the ability of *Legionella pneumophila* to alter eukaryotic host cell migration (57). Beyond bacteria and eukaryotes, recent work demonstrates that quorum-sensing-mediated cross-communication occurs between bacteria and phages (58–60). Specifically, an AI called DPO that is produced by many species of bacteria is detected by a phage that uses the information encoded in DPO to determine whether to enter the lytic or the lysogenic state (59). Our present work provides evidence for a virus with the potential to make MHF. Presumably, inter-domain chemical interactions could enable sharing of and cheating on public goods, enhanced symbiosis or predation, or could allow particular organisms to co-establish niches in otherwise inhospitable environments. Notably, bacteria that synthesize AI-2 generally use it to regulate their own quorum-sensing-controlled behaviors (61). We do not yet know if *S. cerevisiae* and other eukaryotes use MHF as a quorum-sensing signal. Our intention now is to perform RNA-seq on *cff1Δ S. cerevisiae* in the presence and absence of exogenously-supplied MHF to discover the endogenous response.

MHF is a volatile compound that is used as a flavorant. Food scientists have previously shown that low levels of MHF exist in fermented foods, such as soy sauce and malt (62–64). As alluded to above, MHF is hypothesized to form spontaneously from pentose sugars that undergo the Maillard reaction (i.e., during cooking) or as a byproduct (in fungi). Specifically, in the fungus *Zygosaccharomyces rouxii*, it is proposed that, following production of ribulose-5-phosphate by the pentose phosphate pathway, MHF forms spontaneously as a breakdown product (42). Notably, MHF is also a breakdown product of DPD; however, *S. cerevisiae* lacks LuxS, so DPD is an unlikely source of MHF in fungi (44, 65). Importantly, production of MHF by the pentose phosphate pathway and by the Malliard reaction is non-enzymatic. Here, we show that Cff1p, which is presumably an enzyme, is required for MHF production in *S. cerevisiae*. Therefore, if the above non-enzymatic pentose phosphate route to MHF is plausible, there exists an additional enzymatic route involving Cff1p. *Z. rouxii* contains a *CFF1* homolog, suggesting the pathway we have discovered here could also be relevant for *Z. rouxii*.

*S. cerevisiae CFF1* is a gene of unknown function. The crystal structure of *S. cerevisiae* Cff1p has been solved and shows similarity to sugar epimerases and isomerases (26). The Cff1p protein, like other cupins, contains a jelly-roll fold similar to those in germin- and auxin-binding proteins. In Cff1p, the regions adjacent to the cupin motif are distinct from previously studied members of this protein superfamily. Cff1p has notable similarity to the *Salmonella typhimurium* protein, RmlC, which catalyzes the conversion of dTDP-6-deoxy-D-xylo-4-hexulose to dTDP-6-deoxy-L-lyxo-hexulose (66). The crystal structure of RmlC bound to a substrate analog has been compared to that of *S. cerevisiae* Cff1p and shows that Cff1p possesses a binding pocket that could accommodate both the nucleotide with which Cff1p was co-crystallized, and a sugar moiety. However, no sugar was present in the Cff1p crystal. The authors hypothesized that the pocket may bind a sugar-nucleotide. Accordingly, the jelly-roll fold motif is shared with enzymes such as phosphoglucose isomerase and dTDP-4-keto-6-deoxy-D-hexulose-3,5-epimerase (26). Given this relatedness and the fact that DPD, the non-borated precursor to AI-2, is a sugar, we suspect that Cff1p could be the synthase for MHF, and that MHF is made from a sugar substrate.

Our study suggests that the scope of organisms that can participate in quorum sensing through AI-2-type pathways continues to increase, hinting that AI-2 quorum sensing mediates inter-species bacterial communication and inter-domain communication between bacteria and eukaryotes and possibly viruses. Bacteria can distinguish among closely related quorum-sensing AIs and they are capable of decoding and integrating the information contained in blends of AIs to drive appropriate behaviors based on the cell density and the species identities of neighboring bacteria. Co-occurrences of organisms from different domains have important ecological and medical implications (67–70). We highlight a few examples involving *S. cerevisiae*, the main focus of the present work; the presence of *S. cerevisiae* improves the ability of *Pseudomonas putida* to grow in glucose-containing medium (71), enhances the growth of lactic acid bacteria in nitrogen-rich environments (72), and stimulates the growth of multiple *Acinetobacter* species through the secretion of ethanol (73). Our future work will focus on how domain-spanning quorum-sensing cross-communication influences the behavior of the various participants and affects global community structures and their functioning.

## Methods

### Strains, plasmids, and media

Strains and plasmids used in this work are provided in Table S1 and Table S2, respectively. *YML079W (CFF1)* and *YML063W* (*RPS1B*) were deleted from *S. cerevisiae* using the standard kanMX insertion technique (38, 74). Geneticin (G418 sulfate) (Thermo Fisher Scientific) was added at 200 μg/mL for KanMX selection. *S. cerevisiae* was grown in YPD medium, unless specified otherwise (Thermo Fisher Scientific). *V. harveyi* was grown in Luria-Marine (LM) medium and AI-2 reporter assays were performed in AB medium (29). To generate pRS416-*CFF1*, *CFF1* and ~150 bp upstream and downstream were cloned into the XhoI and XbaI (New England Biolabs) sites of pRS416 using standard cloning procedures. We tested two constructs, one containing the *CFF1* gene and 500 bp of upstream and downstream DNA and one containing the *CFF1* gene with upstream and downstream regions of ~150 bp and encompassing only intergenic sequences between *CFF1* and its neighboring genes. Both constructs drove production of the same amount of MHF, suggesting that all of the necessary promoter and terminator elements for *CFF1* are in the intergenic regions. Therefore, we used the construct containing *CFF1* and only the intergenic regions for the work reported here. DNA encoding the *HALO* sequence was fused to that encoding the 3’-terminus of *CFF1* using Gibson Assembly (New England Biolabs) (75). DNA encoding *CFF1* homologs was codon optimized and synthesized (Integrated DNA Technologies) before being inserted between the upstream and downstream regions of *S. cerevisiae CFF1*, cloned into pRS416 (76), and HALO tagged using the strategy outlined above or using Gibson assembly. Mutations in the constructs were generated by standard mutagenic PCR using PfuUltra polymerase (QuikChange II; Agilent Technologies).

### Yeast and mammalian AI-2 mimic production

*S. cerevisiae* cells were grown overnight in YPD or SD-ura (as needed for plasmid maintenance) with shaking at 30 °C. The cells were pelleted by centrifugation for 10 min at 4,000 RPM. The pellet was washed twice with sterile water followed by centrifugation as above. The cells were resuspended in water and incubated for 24 h at 30 °C with shaking. For making yeast AI-2 mimic for molecule identification, the cells were resuspended at OD_600_ = 100 and incubated overnight at 30 °C. When making yeast AI-2 mimic for all other assays, unless otherwise stated, the cells were resuspended at OD_600_ = 10 and incubated overnight at 30 °C. The cells were removed by centrifugation as above, and the resulting cell-free fluids were filtered through 0.22 μm PES membranes (MilliporeSigma). Such preparations were used as the sources of yeast AI-2 mimic for activity assays and for further purification. The mammalian AI-2 mimic produced by Caco-2 cells was prepared as described previously (13).

### V. harveyi TL-25/TL-26 reporter assays

Bioluminescence reporter assays using *V. harveyi* strains TL-25 and TL-26 were performed as previously reported (13, 77). Briefly, vibrio strains were grown overnight at 30 °C in LM medium with shaking. *V. harveyi* cultures were diluted 1:1,000 in AB medium containing 100 μM boric acid. In all cases, cultures were aliquoted into wells of black-sided, clear-bottom 96-well plates (Corning). DPD, AI-1, MHF, or mammalian AI-2 mimic preparation was added at the indicated amounts and the mixtures were serial diluted. Following incubation at 30 °C with shaking for 6 h, bioluminescence and cell density (OD_600_) were measured using an Envision plate reader (PerkinElmer). Data are presented as relative light units (RLU), which are bioluminescence per OD_600_. “Max AI-2” shown in figures indicates bioluminescence output following addition of 125 nM AI-2 (i.e., *S*-THMF-borate). “Max AI-1” depicted in figures indicates bioluminescence output following addition of 500 nM AI-1 (i.e., 3-hydroxyl C4-homoserine lactone).

### Yeast growth curve and survival assays

*S. cerevisiae* strains were grown overnight at 30 °C in YPD medium with shaking to saturation. The cultures were diluted to OD_600_ = 0.1 in a black-sided, clear-bottom 96-well plate. Cells were incubated at 30 °C with shaking and timepoints were taken every 20 min. Growth was measured in a Synergy plate reader (BioTek). For survival analyses, *S. cerevisiae* strains were diluted 1:100,000 and plated on YPD agar plates. Colonies were allowed to grow for 48 h and then counted manually.

### Screen for S. cerevisiae genes required for MHF production

The *S. cerevisiae* Yeast Knockout (YKO) Collection (Dharmacon) is a 96-well plate arrayed library containing about 5,000 *S. cerevisiae* strains with single gene deletions spanning the yeast genome (37–39). After thawing, 5 μL of each strain in the library was transferred into 96-well plates containing 150 μL YPD+G418 sulfate, and the plates were incubated at 30 °C with shaking overnight. 10 μL of each strain was diluted into 150 μL fresh YPD+G418 sulfate medium and allowed to grow for 5 h at 30 °C with shaking. The cells were pelleted at 4,000 RPM for 10 min, and then resuspended in water. The wash and centrifugation steps were repeated two more times. The cells were pelleted at 4,000 RPM for a final 10 min and resuspended in AB medium that had been supplemented with 100 μM boric acid. The initial OD_600_ of each culture was measured to identify any yeast mutants that had grown poorly. Yeast deletion strains that exhibited slowed growth were eliminated from analysis. The plates were incubated overnight at 30 °C with shaking. The following morning, the plates were subjected to centrifugation at 4,000 RPM for 10 min. 75 μL of culture fluid from each well was combined with 75 μL of fresh AB medium that had been inoculated with a 1:1,000 dilution of an overnight culture of the *V. harveyi* TL-26 reporter strain and the mixtures were placed into fresh microtiter plates. The wells were supplemented with 100 μM boric acid (final conc.). Following 8 h incubation at 30 °C with aeration, bioluminescence and OD_600_ were measured (Envision Plate Reader). Normalized RLU were calculated by dividing the bioluminescence by the OD_600_ then dividing by the average RLU from all plates from which measurements were made on a single day. Mutants from the screen that appeared to be defective in production of activity were retested individually, as above. Mutants of interest were reconstructed using the standard KanMX deletion method (38, 74), and again examined for the ability to produce activity.

### Cff1p alignment and phylogenetic tree production

Potential Cff1p homologs were identified using BLASTp (50) and searching against *S. cerevisiae* Cff1p with an E-value cutoff of 1e-5. This analysis returned 1975 sequences. The list of sequences was culled to remove duplicate species resulting in 744 sequences. Rough phylogenetic analyses demonstrated that sequences from species within a genus clustered, thus the list of sequences was further trimmed to include only the top hit in each genus, delivering 367 total sequences. These sequences were aligned using Clustal Omega (78) in SnapGene® software (GSL Biotech). The phylogenetic tree in Figure S7 was constructed in MEGA-X using the Maximum Likelihood method and JTT matrix-based model with 500 bootstrap replications as described previously (79). The alignment was visualized using ggmsa in R, pruning the ends of the alignment to the first and last amino acids of *S. cerevisiae* Cff1p.

### Yeast AI-2 mimic purification and identification

Approximately 250 mL of concentrated crude yeast AI-2 mimic preparation was filtered through a 0.45 μm PVDF filter (MilliporeSigma) and separated on a 2 × 25-cm Luna C18 column (Phenomenex) using a mobile phase consisting of 5% water in methanol at a flow rate of 10 mL/min. The component of interest was eluted in 8 min in approximately 6 mL of mobile phase. The fraction was dried by roto-evaporation to remove methanol. To enable further concentration, the aqueous solution was saturated with sodium chloride and extracted with dichloromethane. The dichloromethane layer containing the product of interest was dried and the sample was refrigerated. Fifty collections were processed in this manner. The products were combined and dissolved in 5 μL of deuterated water to a concentration of 0.003 mg/mL (determined subsequently by HPLC analysis using a standard curve). A ‘background’ sample was generated by collecting the same volume of mobile phase in an area containing no UV visible eluting components. Both samples were analyzed by ^1^H and ^13^C NMR spectroscopy and, although the data were inconclusive, several key signals were detected in the active sample that were not present in and not obscured by the ‘background’ sample. For example, in the ^13^C NMR, signals at δ 194, 172, and 134 ppm were observed reproducibly. These data combined with mass spectrometry data led to a collection of possible candidates for the active component (Fig S2A). The structure was confirmed by GC-MS analysis, including matching of the fragmentation pattern of the active component against a database of known structures. Analysis was carried out on an RTX-1ms (25 μm) 30-m column with 0.32-mm ID (Restek). The samples were incubated at 40° C for 2 min and then subjected to a temperature ramp up of 10° C/min to 300°C, followed by a 5 min hold at 300° C. Mammalian AI-2 mimic activity was separated on a 25 × 0.46 cm Synergi 10 μm Polar-RP column using a mobile phase of 5% water in methanol at a flow rate of 0.5 mL/min.

### Protein Gels

To assess the levels of Cff1p, the Cff1p and Cff homologs, and the Cff1p and Cff variants, yeast cells producing the protein of interest were grown to exponential phase in SD-Ura. In each case, cells equivalent to 10 OD_600_ units were pelleted for 10 min at 4,000 RPM and resuspended in 50 μL Y-PER Yeast Protein Extraction Reagent (Thermo Fisher Scientific) supplemented with 1X Halt Protease Inhibitor Cocktail (Thermo Fisher Scientific). These mixtures were incubated at room temperature for 20 min with agitation and then subjected to centrifugation at 13,000 RPM in a tabletop centrifuge. The supernatants were collected and incubated with 1 μM HaloTag TMR Ligand (Promega) at room temperature for 20 min. 4X Laemmli Sample Buffer was added and the mixtures were incubated at 70 °C for 10 min. Samples were loaded onto 4-20% gradient mini-PROTEAN precast gels (BioRad). Following electrophoresis, proteins were visualized using the Cy3 filter set on an ImageQuant LAS 4000. To assess total protein, gels were stained with Coomassie Brilliant Blue R-250 Staining Solution (BioRad) and visualized using an ImageQuant LAS 4000.

## Acknowledgements

We thank members of the Bassler laboratory for insight and excellent suggestions. We especially thank Jianping Cong who assisted with production of *CFF1* mutant plasmids. *S. cerevisiae* MY8092 and constructs for generating gene deletions were kind gifts of the Rose laboratory. The wild *S. cerevisiae* strains were generous gifts from Dr. Jeffrey Lewis’ laboratory. We also thank Dr. Martin Semmelhack for insight into molecule purification, spectrum interpretation, and molecule identification. We are grateful to Dr. Mohammad Seyedsayamdost and Etienne Gallant for assistance with mass spectral analyses. We are indebted to Valerie Ciraulo at Firmenich who played an essential role in confirming the MHF structure by GC-MS. We also thank Dr. Mohamed Donia for help with constructing phylogenetic trees. This work was supported by the Howard Hughes Medical Institute and NIH grant 5R37GM065859. The content is solely the responsibility of the authors and does not necessarily represent the official views of the National Institutes of Health.

---

**Table S1.**
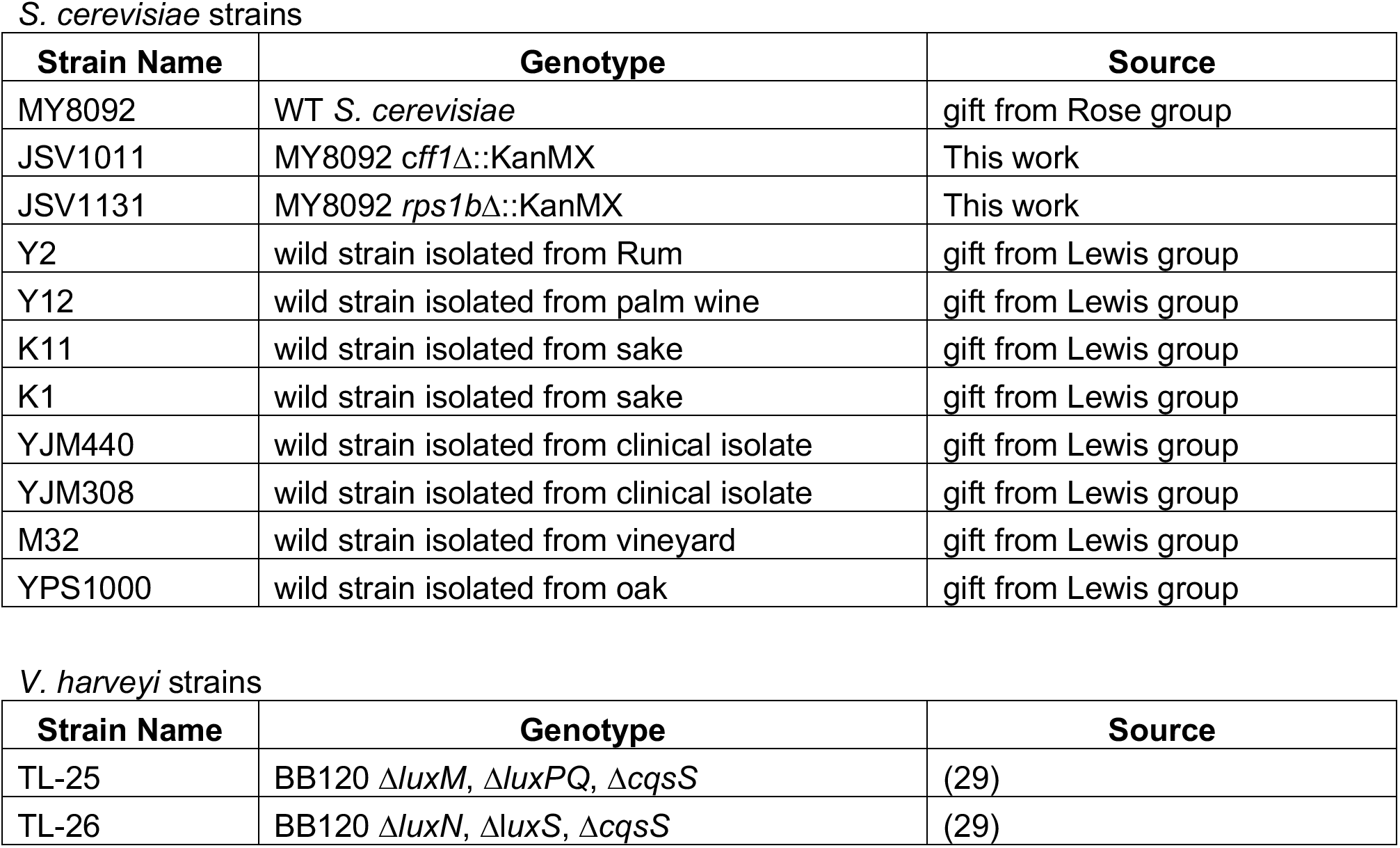
Strains used in this study.

| Strain Name | Genotype | Source |
| --- | --- | --- |
| MY8092 | WT *S. cerevisiae* | gift from Rose group |
| JSV1011 | MY8092 *cff1*Δ::KanMX | This work |
| JSV1131 | MY8092 *rps1b*Δ::KanMX | This work |
| Y2 | wild strain isolated from Rum | gift from Lewis group |
| Y12 | wild strain isolated from palm wine | gift from Lewis group |
| K11 | wild strain isolated from sake | gift from Lewis group |
| K1 | wild strain isolated from sake | gift from Lewis group |
| YJM440 | wild strain isolated from clinical isolate | gift from Lewis group |
| YJM308 | wild strain isolated from clinical isolate | gift from Lewis group |
| M32 | wild strain isolated from vineyard | gift from Lewis group |
| YPS1000 | wild strain isolated from oak | gift from Lewis group |

*V. harveyi* strains
| Strain Name | Genotype | Source |
| --- | --- | --- |
| TL-25 | BB120 $\Delta luxM$, $\Delta luxPQ$, $\Delta cqsS$ | (29) |
| TL-26 | BB120 $\Delta luxN$, $\Delta luxS$, $\Delta cqsS$ | (29) |

**Table S2.** Plasmids used in this study.

| Plasmid Name | Plasmid Description | Source |
| --- | --- | --- |
| pJSV789 | pRS416-p<sub>CFF1</sub>-HALO | This work |
| pJSV751 | pRS416- p<sub>CFF1</sub>-CFF1-HALO | This work |
| pJSV790 | pRS416- p<sub>CFF1</sub>-CFF1-E44A-HALO | This work |
| pJSV784 | pRS416- p<sub>CFF1</sub>-CFF1<sub>*B. cinerea*</sub>-HALO | This work |
| pJSV815 | pRS416- p<sub>CFF1</sub>-CFF1<sub>*B. cinerea*</sub>-E38A-HALO | This work |
| pJSV783 | pRS416- p<sub>CFF1</sub>-CFF1<sub>*T. versicolor*</sub>-HALO | This work |
| pJSV787 | pRS416- p<sub>CFF1</sub>-CFF1<sub>*T. versicolor*</sub>-E30A-HALO | This work |
| pJSV777 | pRS416- p<sub>CFF1</sub>-CFF1<sub>*P. chlororaphis*</sub>-HALO | This work |
| pJSV812 | pRS416- p<sub>CFF1</sub>- CFF<sub>*U. tangerina*</sub>-HALO | This work |
| pJSV809 | pRS416- p<sub>CFF1</sub>- CFF<sub>*S. aureus*</sub>-HALO | This work |
| pJSV806 | pRS416- p<sub>CFF1</sub>- CFF<sub>*A. boritolerans*</sub>-HALO | This work |
| pJSV810 | pRS416- p<sub>CFF1</sub>- CFF1<sub>*A. fumigatus*</sub>-HALO | This work |
| pJSV807 | pRS416- p<sub>CFF1</sub>- CFF1<sub>*C. neoformans*</sub>-HALO | This work |
| pJSV811 | pRS416- p<sub>CFF1</sub>- CFF1<sub>*O. lucimarinus*</sub>-HALO | This work |
| pJSV813 | pRS416- p<sub>CFF1</sub>- CFF1<sub>*S. kowalevskii*</sub>-HALO | This work |
| pJSV805 | pRS416- p<sub>CFF1</sub>- CFF1<sub>*M. euhalobius*</sub>-HALO | This work |
| pJSV808 | pRS416- p<sub>CFF1</sub>- CFF1<sub>*A. hypogyna*</sub>-HALO | This work |
| pJSV814 | pRS416- p<sub>CFF1</sub>- CFF1<sub>*P. salinus*</sub>-HALO | This work |

**Figure S1.**
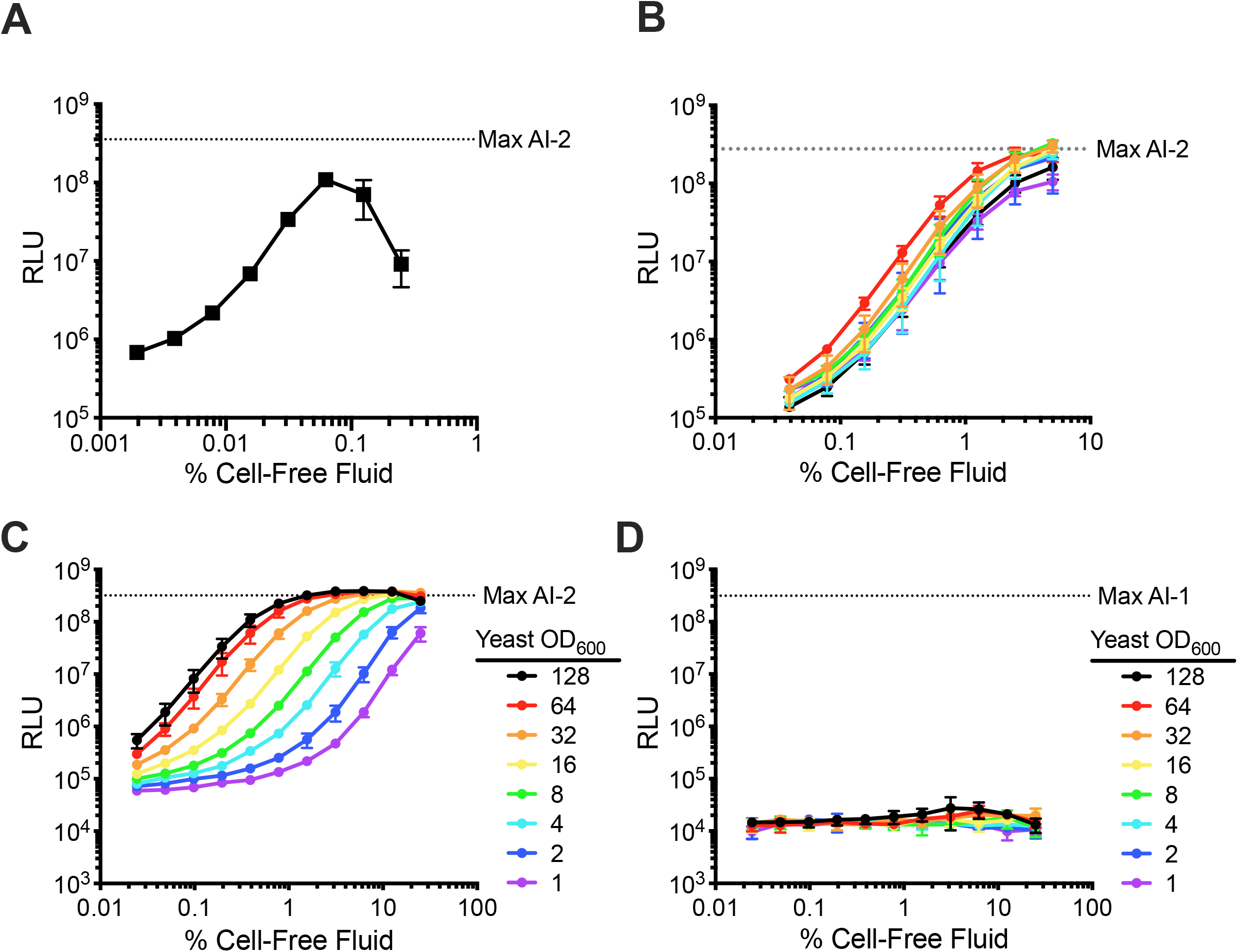
*S. cerevisiae* isolates produce an AI-2 mimic. (A) Light output from the *V. harveyi* TL-26 reporter strain in response to cell-free fluids from *S. cerevisiae* grown for 48 h in synthetic defined medium with glucose as the carbon source. (B) Light output from the *V. harveyi* TL-26 reporter strain in response to cell-free fluids from wild *S. cerevisiae* isolates grown to OD_600_ = 25. The colors represent the *S. cerevisiae* isolates: red, K1; orange, YJM308; yellow, YJM440; green, M32; cyan, YPS1000; blue, Y12; purple, Y2; black, K11. (C) Light output from the *V. harveyi* TL-26 reporter strain in response to cell-free fluids from *S. cerevisiae* grown to the designated OD_600_ values. (D) As in panel C, with the *V. harveyi* AI-1 reporter strain TL-25. RLU and Max AI-2 as in Figure 1. Max AI-1 denotes activity from 500 nM AI-1. In all panels, error bars represent standard deviations of biological replicates, N=3.

**Figure S2.**
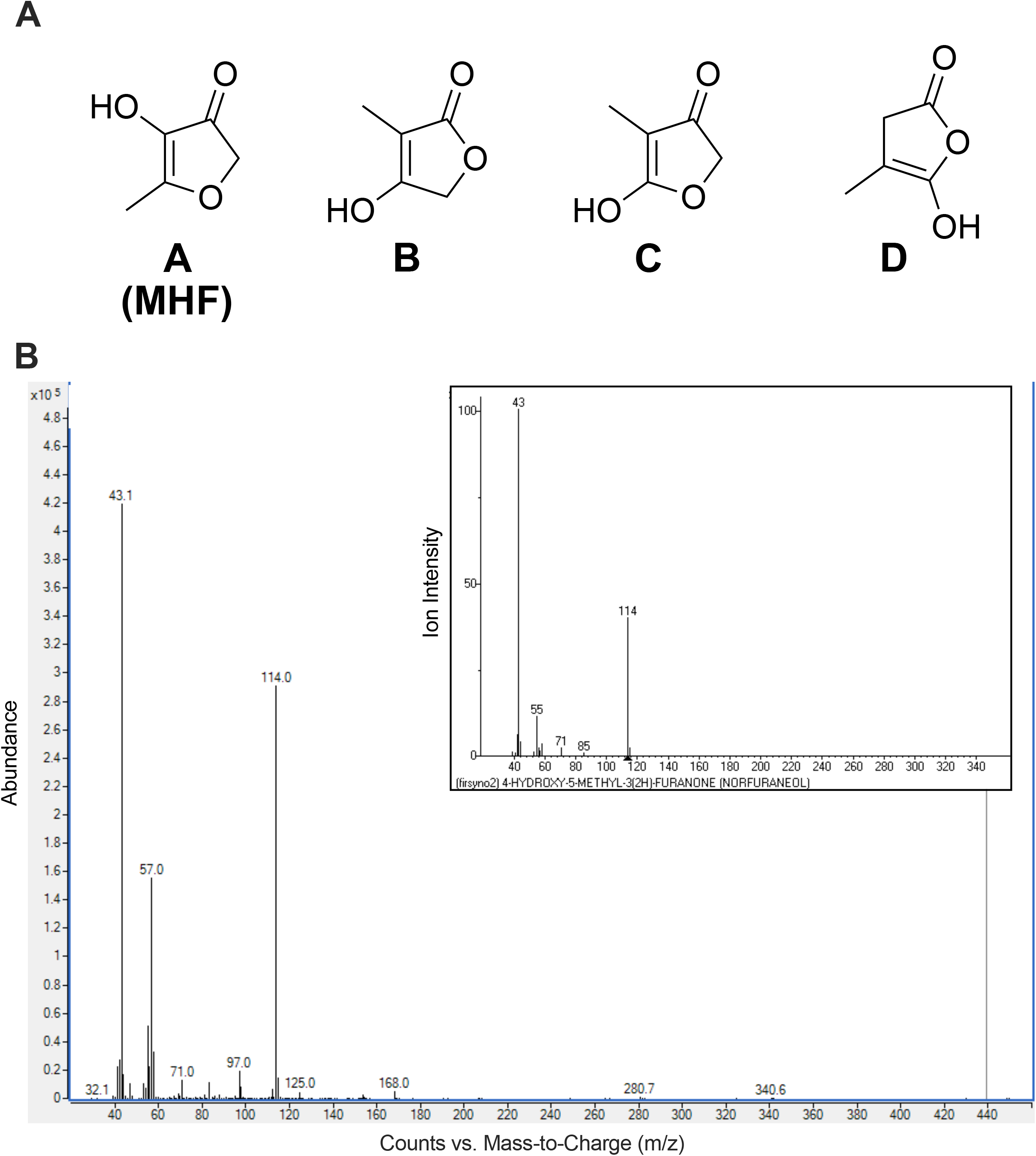
Identification of the AI-2 mimic as MHF. (A) Compounds **A**-**D** proposed as candidate structures for the yeast AI-2 mimic. (B) GC-MS profile for the peak containing the active component from the yeast AI-2 mimic preparation. Inset, National Institute of Standards and Technology (NIST) database GC-MS spectrum for MHF.

**Figure S3.**
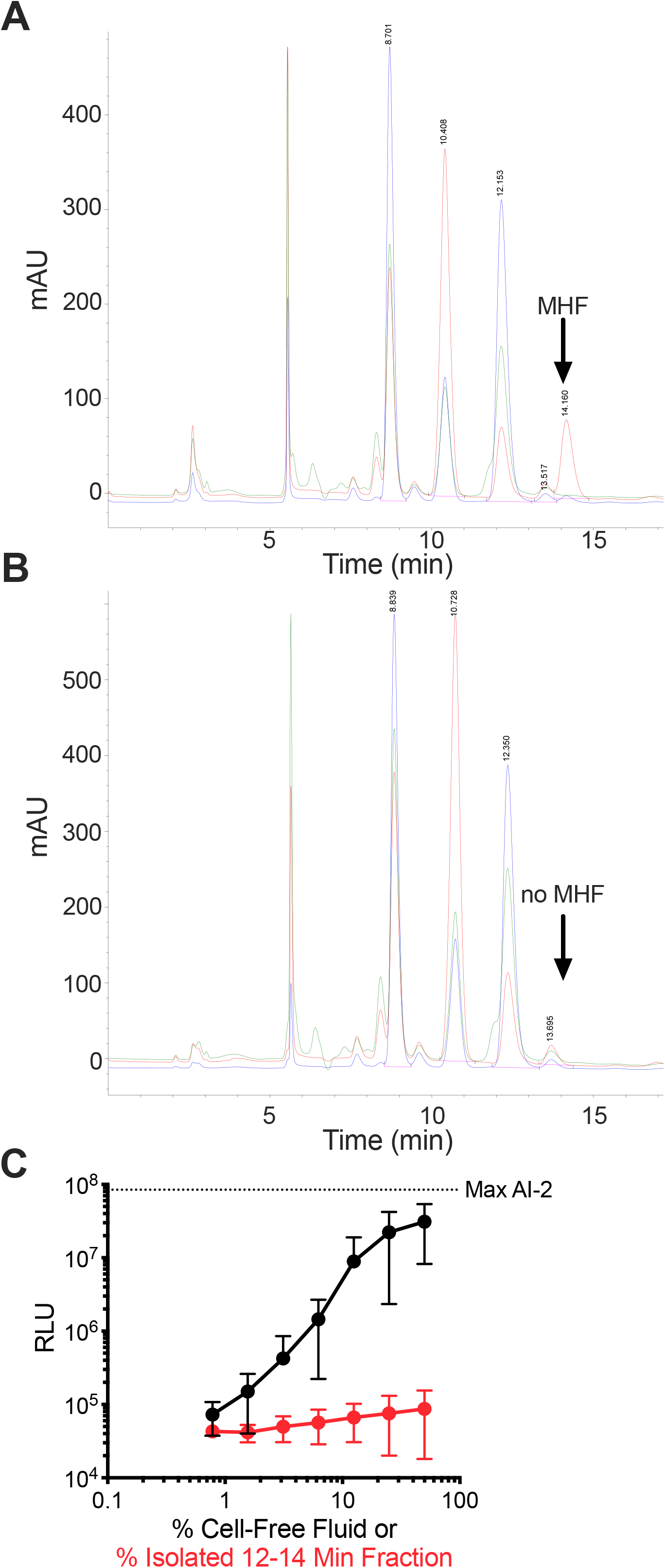
The mammalian AI-2 mimic is not MHF. (A) The mammalian AI-2 mimic preparation was fractionated by HPLC with a spike of 2 μg/mL of commercial MHF (arrow). (B) As in (A) but lacking the MHF spike. The arrow shows where MHF would elute if present. (C) Light output from the *V. harveyi* TL-26 reporter strain in response to the mammalian AI-2 mimic prior to HPLC fractionation (black) and activity from the 12-14 min fraction (red). In C, error bars represent standard deviations of biological replicates, N=3. mAU = milli-Absorbance Units. RLU and Max AI-2 as in Figure 1.

**Figure S4.**
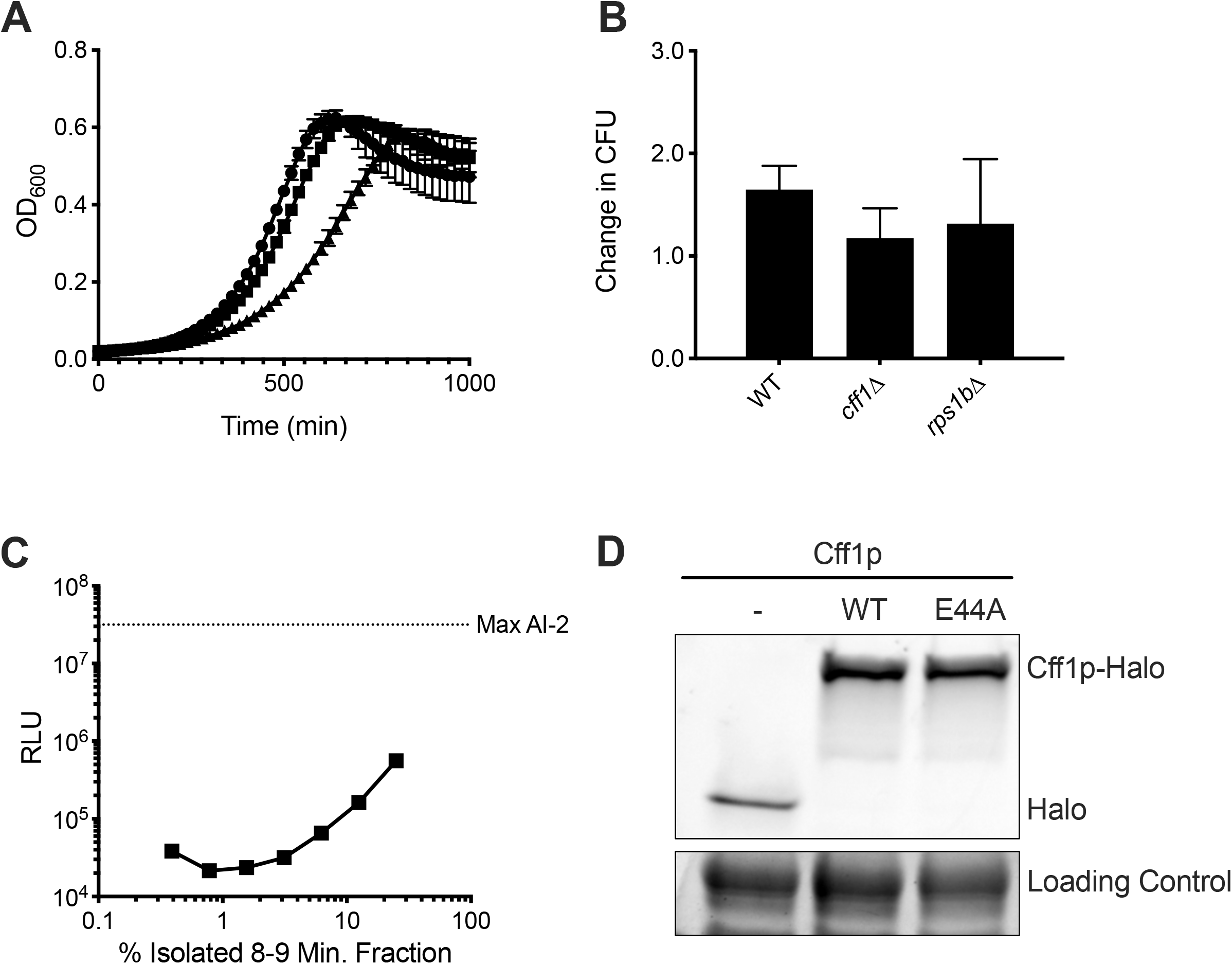
Growth and survival of *S. cerevisiae* mutants, their MHF production profiles, and Cff1p abundance. (A) Growth curves of WT *S. cerevisiae* (squares), *cff1Δ S. cerevisiae* (circles), and *rps1bΔ S. cerevisiae* (triangles). OD_600_ was monitored every 20 min. (B) Survival of the strains in A following overnight incubation in water at 30 °C. Change in CFU (Colony Forming Units) is calculated as CFU after incubation divided by CFU before incubation. (C) Light output by the *V. harveyi* TL-26 reporter strain in response to activity present in the 8-9 min HPLC fraction from an AI-2 mimic preparation made from *cff1Δ S. cerevisiae.* (D) Levels of the designated Cff1p-HALO proteins visualized by staining with HaloTag TMR ligand. The loading control shows a prominent band from Coomassie staining of the gel. RLU and Max AI-2 as in Figure 1. In A, B, and C, error bars represent standard deviations of biological replicates, N=3.

**Figure S5.**
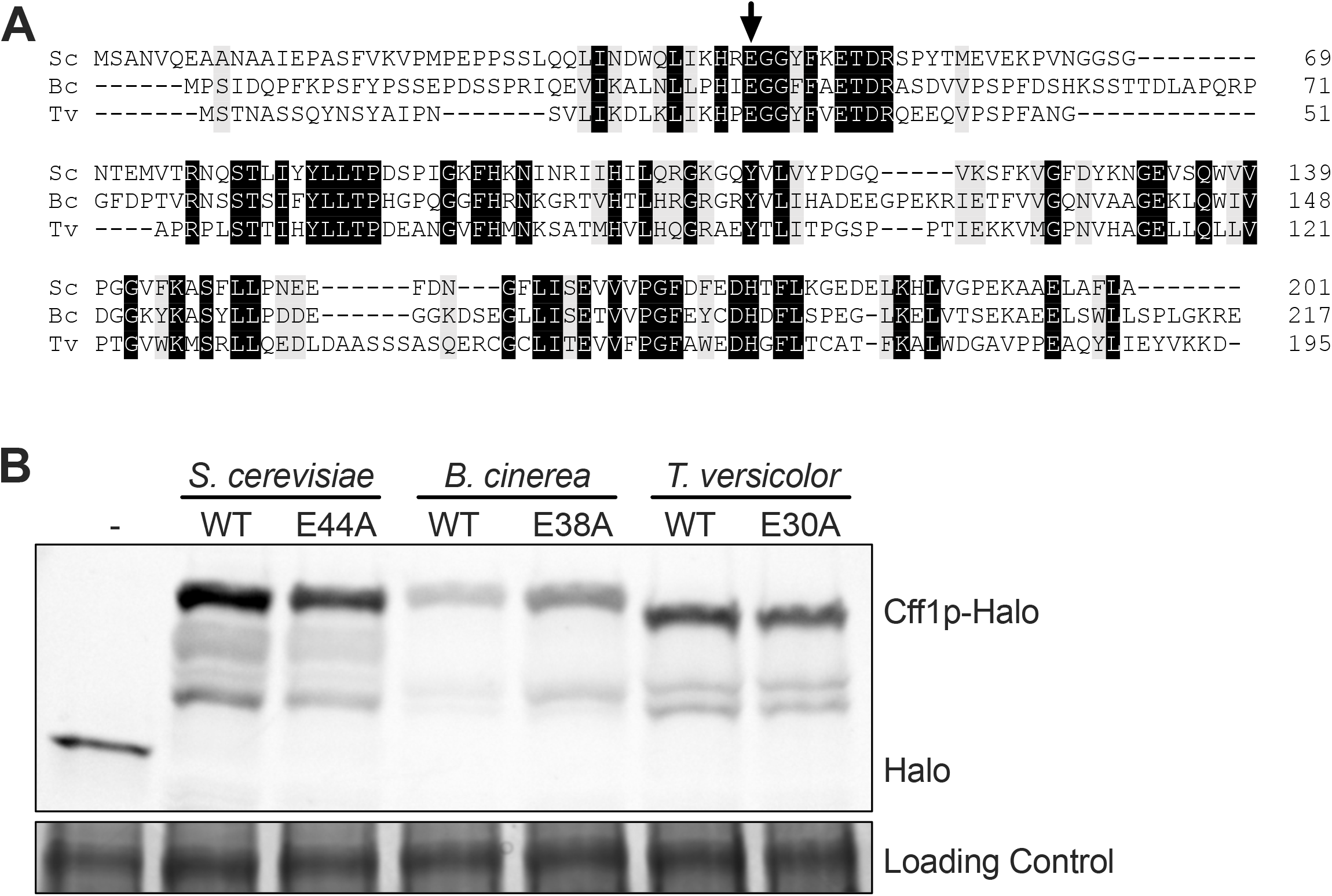
Comparison of Cff1p homologs and their abundances in *cff1Δ S. cerevisiae*. (A) Amino acid sequence alignment of Cff1p homologs from *S. cerevisiae* (Sc), *B. cinerea* (Bc), and *T. versicolor* (Tv). Black and gray boxes designate amino acid identity and similarity, respectively. The residues equivalent to E44 in *S. cerevisiae* are shown by the arow (B) Protein gel showing levels of WT and mutant Cff1p-Halo from *S. cerevisiae*, *B. cinerea*, and *T. versicolor* produced by *cff1Δ S. cerevisiae* carrying the corresponding genes. Cff1p-Halo and Halo bands are labeled. Additional bands, likely protein degradation products, are present. The loading control is a prominent band from Coomassie staining of the gel.

**Figure S6.**
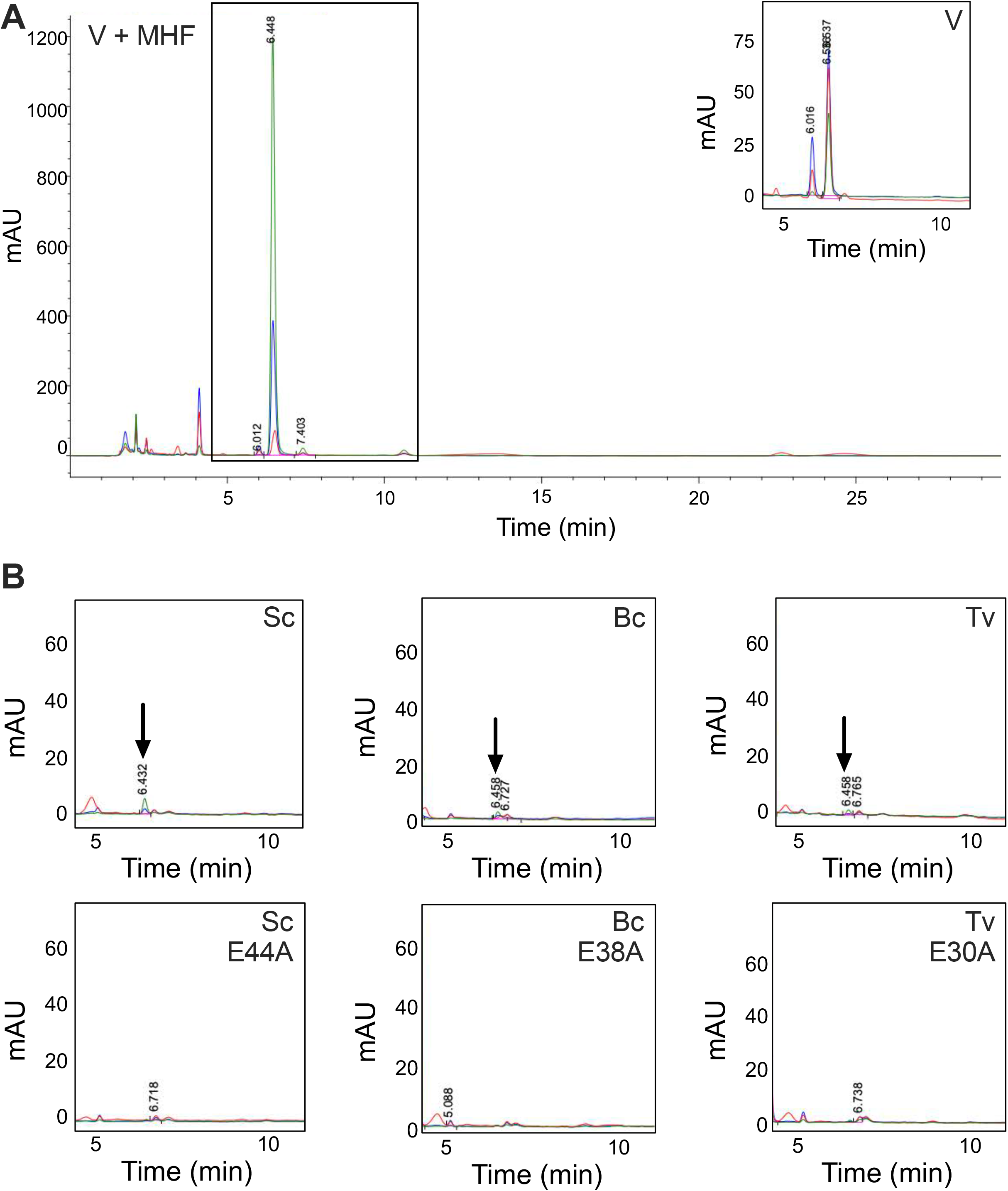
Homologs of Cff1p can complement the loss of MHF production in *cff1Δ S. cerevisiae*. (A) Chromatogram depicting HPLC fractionation of yeast AI-2 mimic prepared from *cff1Δ S. cerevisiae* carrying an empty vector (denoted V) (inset) or supplemented with 125 ng exogenous MHF. The chromatograms show absorption at 214 (red), 254 (blue), and 280 (green) nm. The boxed region shows that MHF absorbs at 254 (blue) and 280 (green) nm and elutes at 6.448 min. This region should be compared to that in the inset showing that this peak is absent in the same sample that did not receive exogenous MHF. (B) Segments of chromatograms showing the same regions boxed in panel A for fractionations of yeast AI-2 mimic prepared from *cff1Δ S. cerevisiae* carrying WT and the designated mutant *CFF1* alleles from *S. cerevisiae* (Sc), *B. cinerea* (Bc), and *T. versicolor* (Tv). The arrows in the top row of boxes denote the MHF peaks in the samples prepared from *S. cerevisiae* carrying each WT *CFF1* gene. The bottom row of boxes show that those peaks are absent when *S. cerevisiae* carries the mutant genes. In all panels, absorption at 214 nm is shown in red, 254 nm is shown in blue, and at 280 nm is shown in green. Note that the color scheme differs from other figures with HPLC traces. mAU = milli-Absorbance Units.

**Figure S7.**
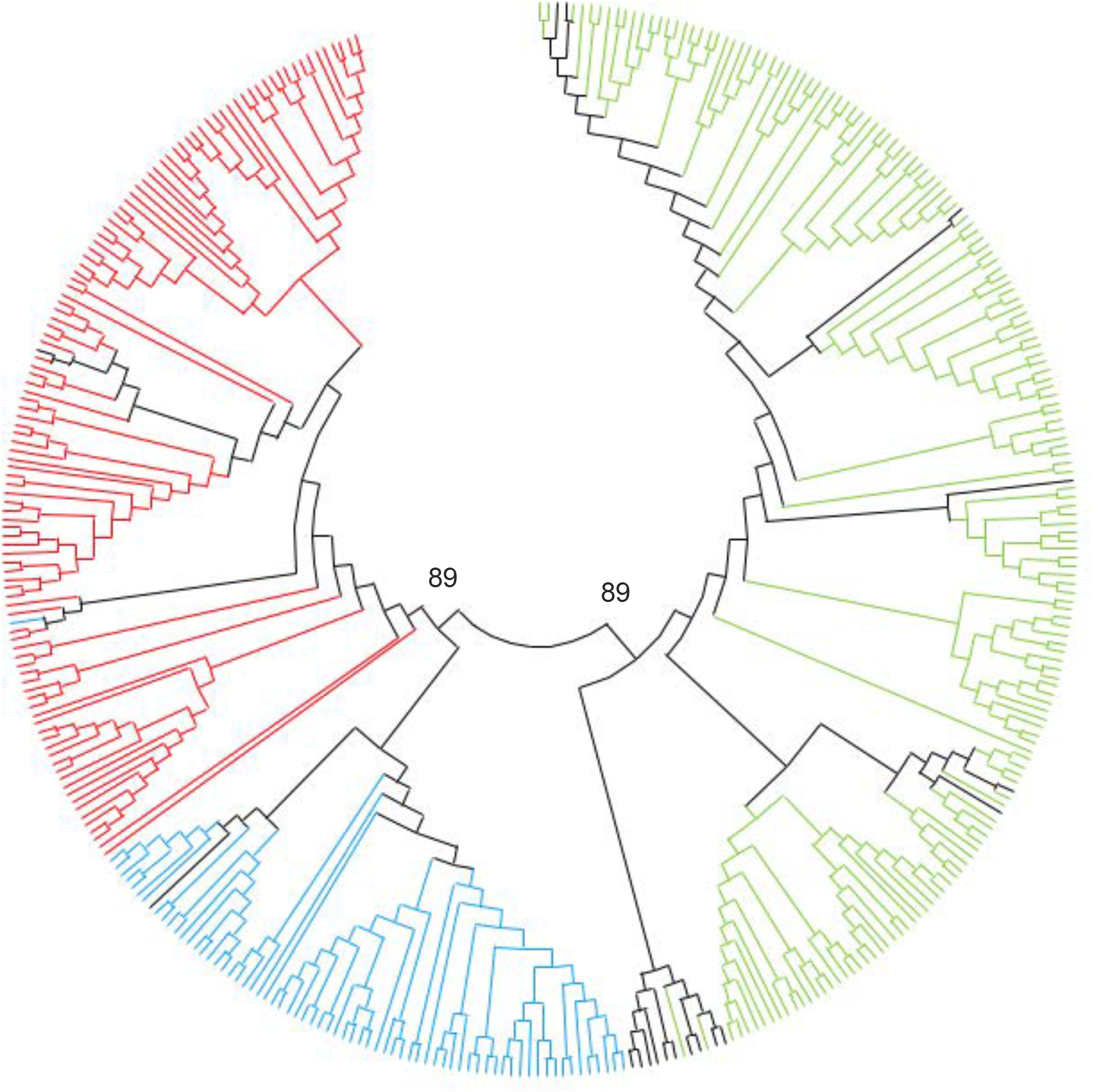
Organisms from all domains contain Cff1p homologs – expanded analysis. Phylogenetic tree showing all genera with Cff1p homologs uncovered by BLASTp analysis (50). Red, fungi species belonging to the Ascomycota phylum; blue, fungi species belonging to the Basidiomycota phylum; green, bacterial species; black, all other species. The frequencies of the two highest level nodes, following 500 bootstrap replications, are shown.

